# pyALRA: python implementation of low-rank zero-preserving approximation of single cell RNA-seq

**DOI:** 10.1101/2025.03.20.644345

**Authors:** Alexandre Lanau, Joshua J Waterfall

## Abstract

**Summary:** We present pyALRA, an efficient python implementation of the ALRA R package conceived to impute drop out values using a low-rank zero-preserving approximation for single cell RNA-seq. This re-implementation achieves similar performance prediction using corresponding python methods and allows both speed and RAM consumption improvements.

**Availability:** pyALRA is released as an open-source software under the MIT license. The source code is available on GitHub at *https://github.com/alexandrelanau/pyALRA*.

## INTRODUCTION

Single-cell RNA sequencing (scRNA-seq) has revolutionized transcriptomic research by enabling the measurement of gene expression at the individual cell level, providing invaluable insights into cellular heterogeneity and gene regulation. However, a prominent challenge in analyzing scRNA-seq data is the high prevalence of false zeros or “dropout” events, where genes that are actually expressed may appear as unexpressed due to undersampling or other technical limitations. These dropout events can significantly impede downstream analyses, such as clustering and differential expression, necessitating robust imputation methods that can accurately recover the underlying biological signal while preserving the true biological zeros (genes not expressed at the time point of library generation).

Various approaches have been developed to address this issue, including deep learning-based methods like DCA^1^, graph-based techniques such as MAGIC^2^, and statistical models like SAVER^3^. While these methods have shown improvements in imputing technical zeros, many fail to preserve true biological zeros, potentially leading to overestimation of gene expression across cell populations^4^. ALRA (Adaptively Thresholded Low-Rank Approximation)^4^ was introduced to overcome these limitations by selectively imputing technical zeros based on a low-rank matrix approximation, restoring biological zeros through adaptive thresholding.

However, many of these algorithms remain unavailable in Python, as R has historically been the dominant language in bioinformatics over the past decades. With the recent development of the scverse ecosystem^5^, Python is increasingly becoming the preferred language at the intersection of various bioinformatics domains. Existing methods for interfacing R and Python, such as rpy2 (https://rpy2.github.io/), are often time-consuming and require significant effort, while offering limited control over the interaction between the two languages.

Here, we present a Python implementation of the ALRA methodology for scRNA-seq data imputation, aiming to make this tool accessible to the Python-based bioinformatics community. Our implementation facilitates integration with modern Python data science frameworks, enhances computational scalability, and provides results consistent with the original method. Through benchmarking on multiple datasets, we demonstrate that our implementation achieves comparable performance to existing methods while preserving biological zeros, thus improving the reliability of downstream scRNA-seq analyses. (pyALRA and ALRA are developed by independent groups).

## IMPLEMENTATION

We applied the Adaptively Thresholded Low-Rank Approximation (ALRA) method to several single-cell RNA sequencing (scRNA-seq) datasets to evaluate its performance in imputing expression values while preserving biological zeros. In this implementation, we leveraged only well-maintained packages: numpy, scipy and scikit-learn.

### 2.1 Available features and code structure

ALRA normalizes the input expression matrix by library size and applies a log transformation to stabilize variance, tailoring normalization to the data type. It employs the `randomized_svd` function from the `sklearn.utils.extmath` module to perform randomized singular value decomposition (SVD), adaptively determining the rank of the low-rank approximation based on singular value differences. As described in the original ALRA publication, the rank k that represents biological signals from the SVD is identified as the lowest singular value (SV) which is larger than the subsequent SV by an amount significantly greater than the gaps between SVs in the tail of the distribution (by default lowest 20 SVs). The process reconstructs the matrix and applies a quantile-based thresholding method to impute missing values while preserving biological zeros, enhancing the reliability of downstream analyses.

### 2.2 Benchmark of pyALRA vs ALRA package

To evaluate the relevance of our implementation, we measured prediction performance and computational resource consumption. All predictions are based on the same input matrices, which were previously normalized using the same method. As qualitative evidence, we used the PBMC dataset from ALRA tutorials to compute umap and leiden clustering based on normalized, r-ALRA imputed or pyALRA imputed counts **(Fig.1A)**. r-ALRA and pyALRA presents qualitivatively similar shape and clustering (with regards to seed variation in UMAP). Using alluvial plot, we can identify high sharing of cluster composition between r-ALRA and pyALRA imputed clusters **(Fig.1B)**.

For each dataset, 3 subsets of increasing size were randomly subsampled (1000, 10000 and 50000 cells). And for each subset, 5 independent runs were performed.

To perform quantitative comparisons, when comparing the predicted k values from the randomized SVD method, ALRA and pyALRA present similar predicted k values (**Fig. 1C, S1-S3**). To ensure stability of k predictions, due to the increased efficiency of the randomized SVD used here, we increased the number of iterations (n=10 by default compared to 2 in the original publication). Moreover, when comparing the prediction of non-zero genes by ALRA and pyALRA, both methods show similar percentages of predicted non-zero genes across datasets **(Fig.1D, S1-S3)**.

**Figure 1:**
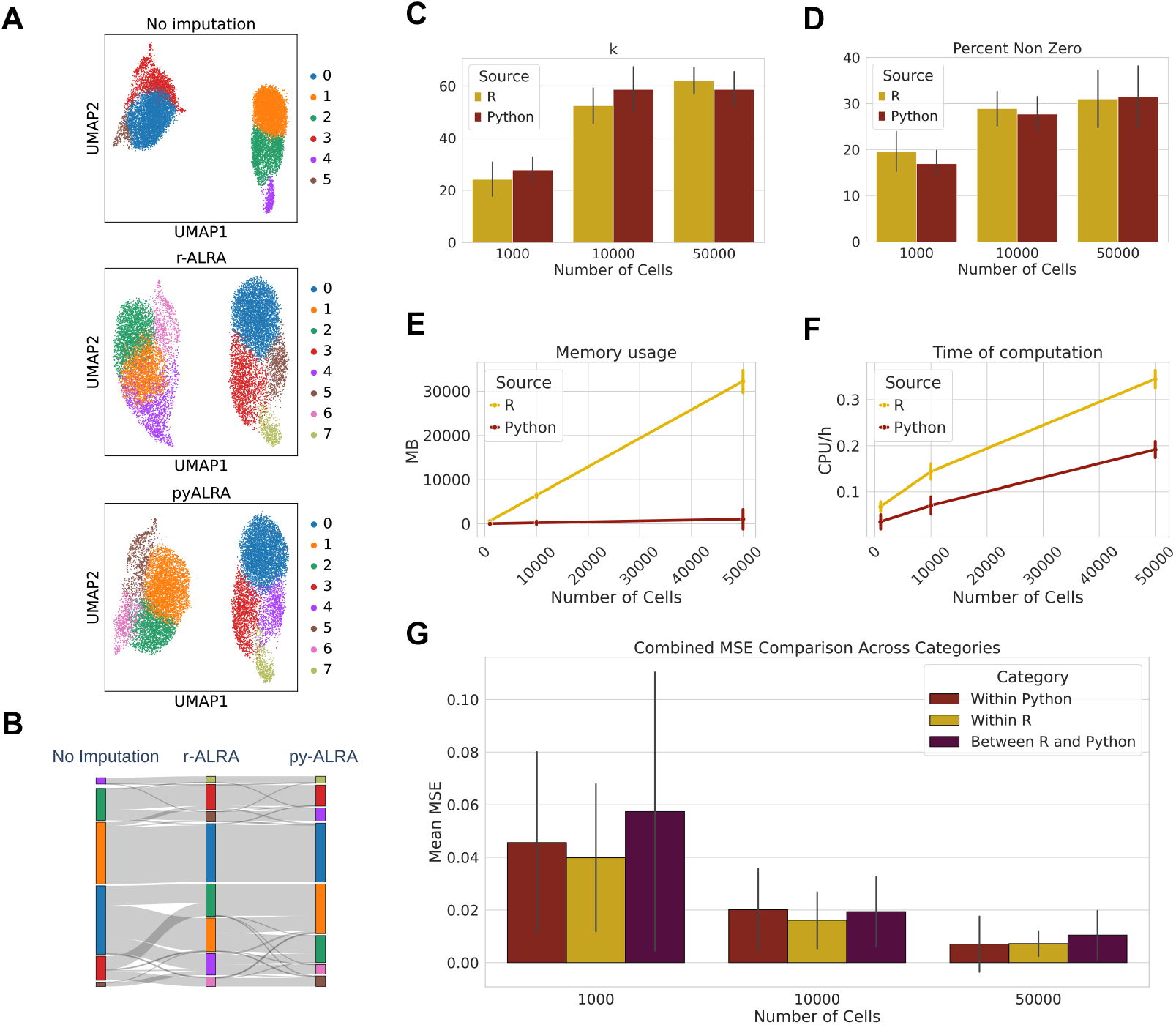
pyALRA presents similar predictions with better computation performances. **(A)** UMAP of single cell RNA sequencing from ALRA PBMC tutorials, only normalized (top panel), r-ALRA processed (middle panel) or pyALRA processed (bottom panel). **(B)** Alluvial plot of cluster composition across the different imputation conditions from (A). **(C)** Comparison of k predicted using randomized SVD algorithm between R and Python implementation (n=15, error bars = standard deviation). **(D)** Comparison of non-zeros genes predicted between R and Python implementation (n=15, error bars = standard deviation). Comparison of Python and R implementation of ALRA for RAM usage (Mb) (**E)** and CPU/h (**F)** (n=15, error bars = standard deviation). **(G)** Mean-squared error comparison between pyALRA and ALRA implementation (n=75 per condition, error bars = standard deviation). For **(C-G)**, all results are from 5 runs in subsamples (1000, 10000, 50000 cells) from 3 independent datasets.

Additionally, we compared the computational performance between the R and Python implementations of the algorithm. PyALRA consistently demonstrates better computational performance, with lower RAM usage and CPU hours consumed (**Fig. 1E-F, S1-S3**).

Finally, by performing mean-squared error of predicted counts between each run for each datasets, within python predicted, within R predicted and between both implementations, we observed a low mse between differents predictions **(Fig.1G, S4)**. As shown in Fig. 1, we observed similar predictions with low mean squared error across the datasets, along with improved computational performance from pyALRA.

### 2.3 Conclusion

In conclusion, pyALRA is an efficient and reliable method to impute technical zeros in single-cell RNA-seq data. It produces equivalent output to the original R implementation with significant improvements in time and memory usage.

## Data availability

The data underlying this article were downloaded from EBI Single Cell Expression atlas. (www.ebi.ac.uk/gca/sc/experiments), using scanpy.datasets.ebi_expression_atlas function from scanpy. Following datasets were used: E-MTAB-8142, E-MTAB-7407, E-GEOD-139324.

## Author contribution

AL and JJW conceived the study and wrote the manuscript. AL wrote the code and performed all analyses under supervision of JJW.

