## Supplemental figures for "pyALRA: python implementation of low-rank zero-preserving approximation of single cell RNA-seq"

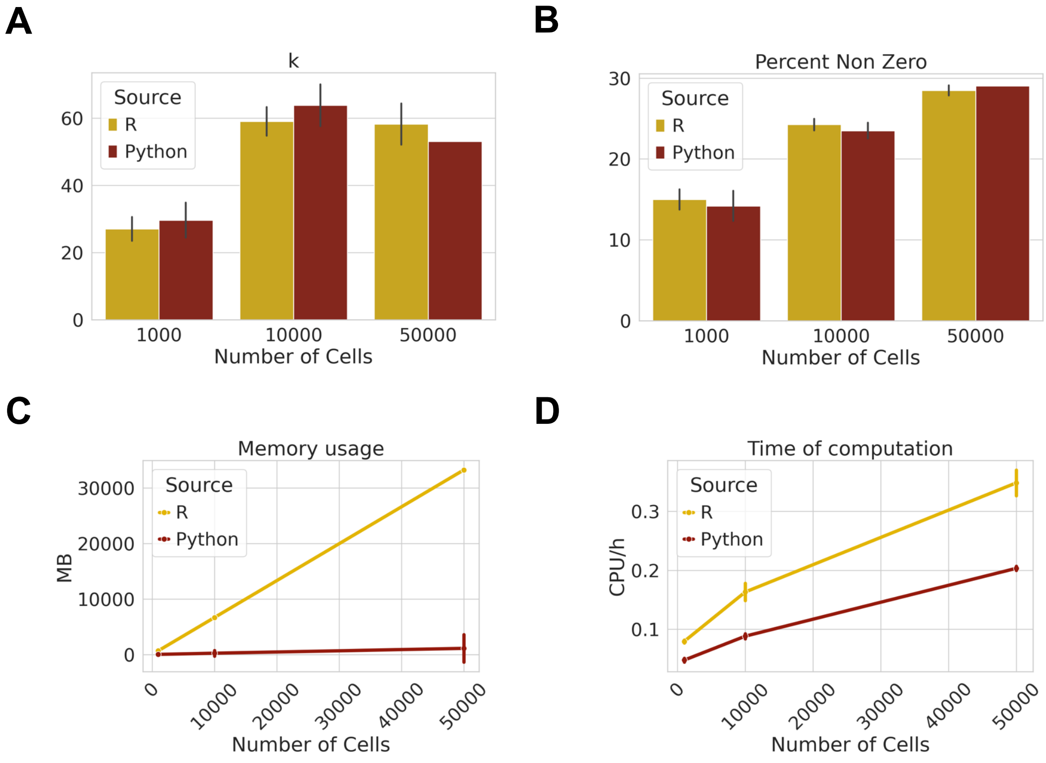


**Figure S1: Performance of prediction for E-MTAB-8142**

(**A**) Comparison of k predicted using randomized SVD algorithm between R and Python implementation (n=15, error bars = standard deviation). **(B)** Comparison of non-zeros genes predicted between R and Python implementation (n=15, error bars = standard deviation). Comparison of Python and R implementation of ALRA for RAM usage (Mb) (**C**) and CPU/h **(D)** (n=15, error bars = standard deviation).

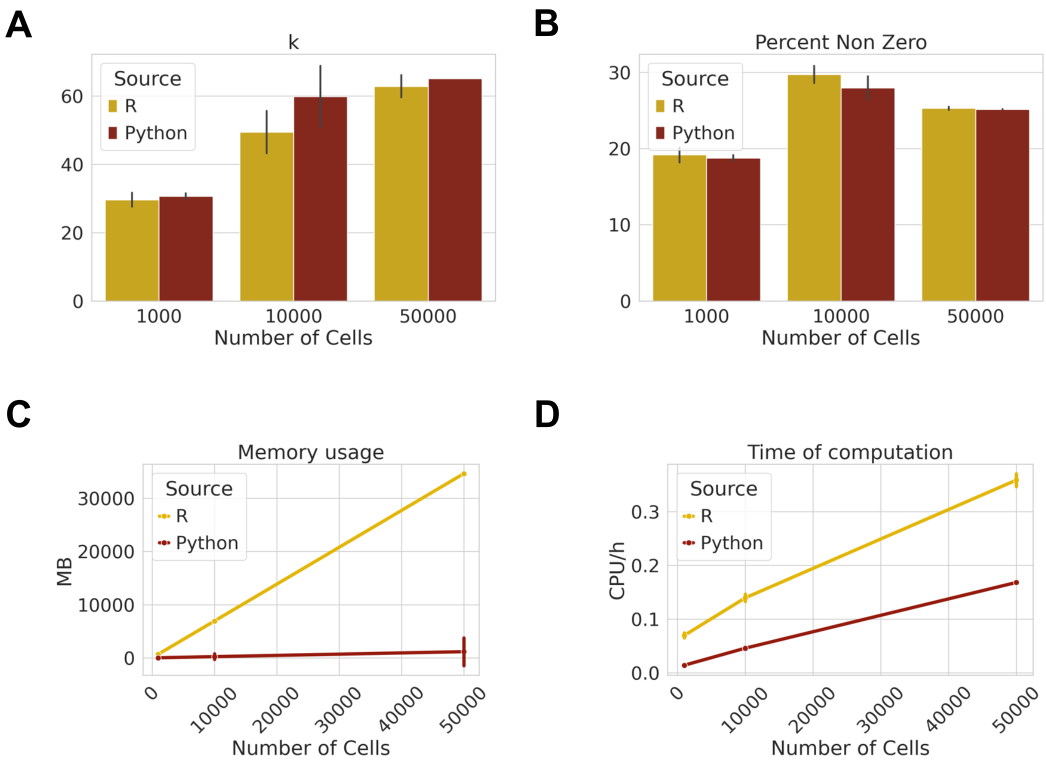


**Figure S2: Performance of prediction for E-MTAB-7407**

(**A**) Comparison of k predicted using randomized SVD algorithm between R and Python implementation (n=15, error bars = standard deviation). **(B)** Comparison of non-zeros genes predicted between R and Python implementation (n=15, error bars = standard deviation). Comparison of Python and R implementation of ALRA for RAM usage (Mb) (**C**) and CPU/h **(D)** (n=15, error bars = standard deviation).

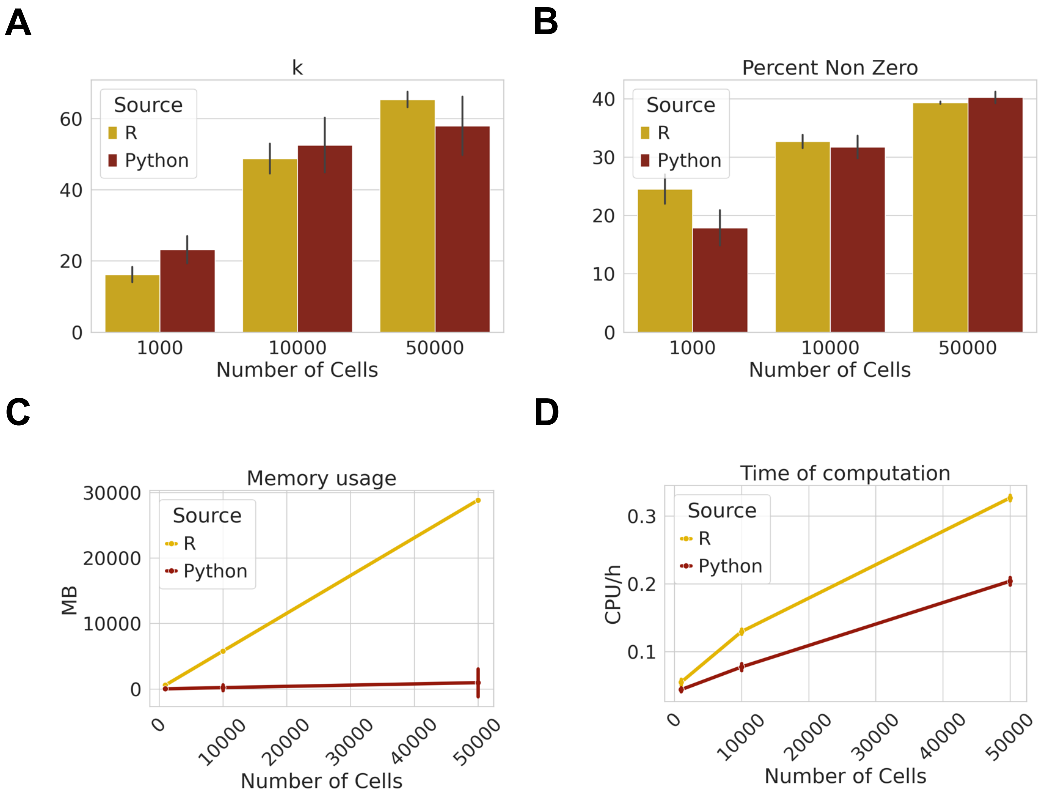


**Figure S3: Performance of prediction for E-GEOD-139324**

(**A**) Comparison of k predicted using randomized SVD algorithm between R and Python implementation (n=15, error bars = standard deviation). **(B)** Comparison of non-zeros genes predicted between R and Python implementation (n=15, error bars = standard deviation). Comparison of Python and R implementation of ALRA for RAM usage (Mb) (**C**) and CPU/h **(D)** (n=15, error bars = standard deviation).

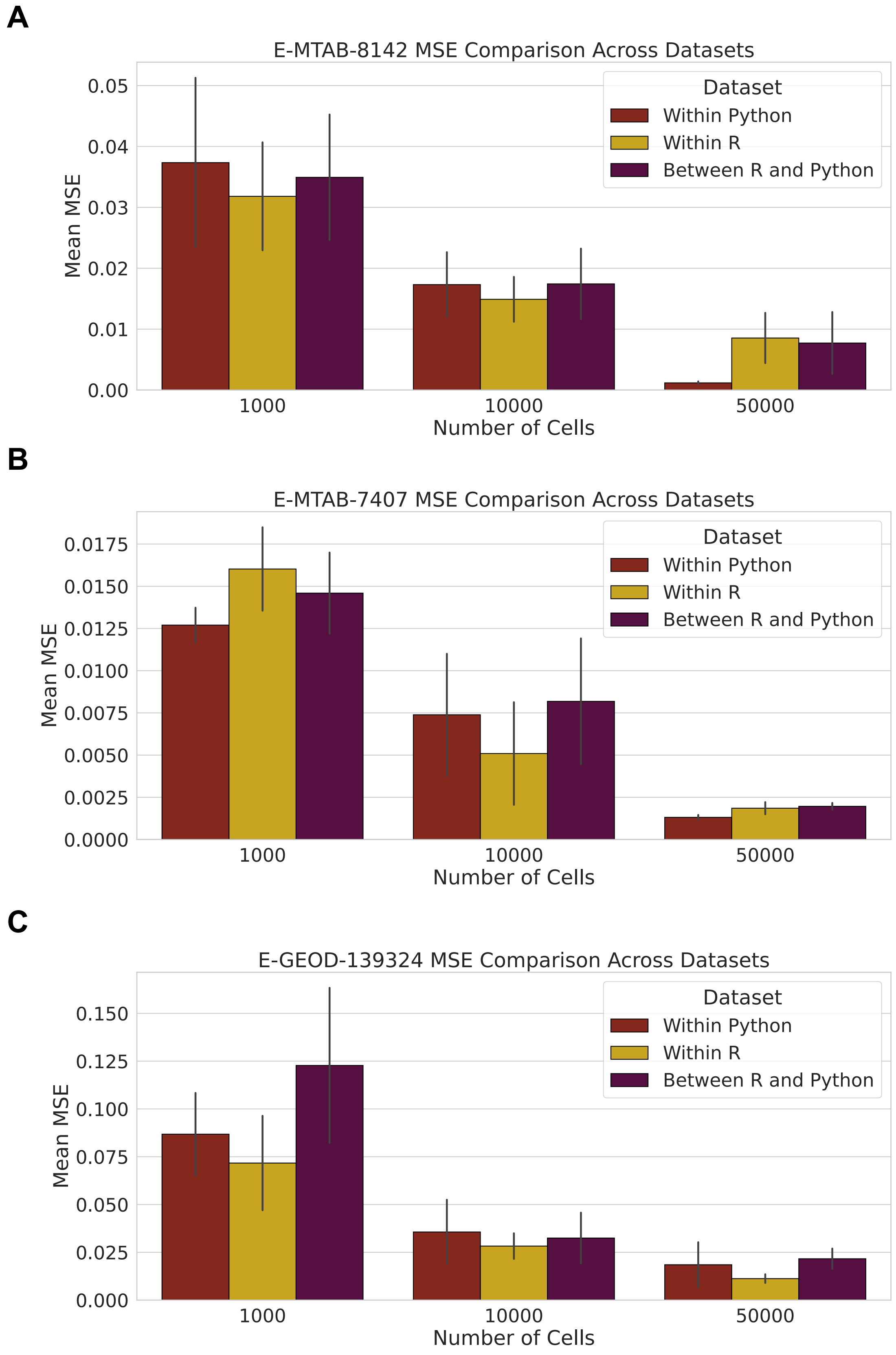


**Figure S4: Pairwise comparisons between pyALRA and r-ALRA runs for E-MTAB-8142, E-MTAB-7407, E-GEOD-139324**

(**A-C)** Pairwise comparison within (R and between R and Python ALRA implementation (right column), and within each implementation (python – left column, R – middle column) , using mean-squared error, for several dataset sizes - 1000 **(A)**, 10000 **(B)** and 50000 cells **(C)** - and between each run.
